# Education influences knowledge about environmental issues in Washington, DC, USA

**DOI:** 10.1101/793810

**Authors:** Matthew L. Richardson, Ashley D. Milton, Elgloria Harrison

## Abstract

We predicted that demographic differences in Washington, DC’s population would influence people’s knowledge and perceptions about the interconnectedness of natural resources, climate change, economics, and socio-cultural well-being, so we conducted surveys over three years to test that prediction. We collected demographic data from 455 participants and asked them 26 questions/statements related to natural resources, climate change, economics, and health. We selected education as the focal demographic category and participants were categorized based on their level of educational attainment: 1) completion of high school or less (hereafter “high school”); 2) some trade school or university education beyond high school up to and including completion of a trade school, two-, or four-year degree (hereafter “post-high school”); and 3) completion of a Master’s, professional, or doctoral degree (hereafter “advanced education”). Answers to 14 of the 26 survey questions were dissimilar across educational groups. People with advanced education reported the highest connection with the natural community and were more likely to report that their personal welfare depended on the natural community. Participants in the high school group were more likely to believe that humans do not have much influence on natural resources and placed more trust in technology and human achievements to control nature and ensure that earth will not become unlivable. Compared to those with education beyond high school, those with a high school education were more likely to express an interest in local environmental concerns over global, jobs over natural resources, and effects of degraded local natural resources on income, health, and the environment instead of on cultural/social practices, neighborhood aesthetics, and recreation. The results suggest ways in which educational information and engagement in environmental issues should be targeted for stakeholders of different educational background in order to increase knowledge and build effective partnerships that find solutions for environmental problems.

## INTRODUCTION

Only 14% of the world’s population lived in cities in 1900, but now over 50% live in cities and this percentage is expected to reach 66% by 2050 [1]. The United States of America (USA) has an even higher urban population than the world average: over 80% live in urban areas [2]. The urbanization of the human population is happening simultaneously with worsening local and global environmental problems, such as overexploitation and degradation of natural resources [3–4], population declines and extinctions of other species [5–6], and climate change [7]. These environmental problems are interrelated in often complex ways and have the potential to influence neighborhood aesthetics and a person’s economic well-being, health, cultural and social practices, and recreation [8–9]. Whereas environmental knowledge does not necessarily lead people to take pro-environmental actions, tackling environmental, and related economic, social, and cultural, problems may be more challenging if the general public is under- or uneducated about the problems [10–12].

Cities can have a profound influence on natural resources and pollution within a region as well as globally, which in turn can negatively affect human well-being [13–15]. Therefore, effective solutions for sustainably using natural resources, curtailing climate change, and improving the lives of people must consider the role that cities can play [8, 16–17]. Some city governments have been more proactive than others in addressing environmental problems and the well-being of the city’s inhabitants. For example, London (UK) and Beijing (China) have made efforts to electrify transportation, including public buses and taxis, in order to improve air quality [18] and Portland, Oregon is considered one of the most advanced cities in the USA for climate planning because they have been conducting work on mitigation since the 1990s [19]. Characteristics of proactive cities include a political culture that embraces mitigation, a general public that has an awareness of environmental problems and advocates that their political leaders act, and local experts that engage with government agencies [19].

Washington, DC is the capital of the USA and its District government has plans for sustainability, improving air quality, adapting to climate change, reducing the government’s carbon footprint, and protecting wildlife and watersheds [20–21]. The District government also commissioned a study on the linkage between urban heat islands and poor health [22]. These plans and studies may indicate that Washington, DC has the characteristics of a proactive city because they seek to identify and mitigate environmental problems and the associated impacts on people. However, little is known about environmental knowledge and perceptions of residents in the Washington, DC area. Globally, environmental knowledge and action are often correlated with demographic characteristics, such as education, age, gender, and place of residence[23–26], and participation in group organizations [12].

Washington, DC has an extremely diverse population, there are large disparities among the population in education, employment, income, health, and overall well-being [27], and distribution of natural, manmade, and financial resources is unequal [21, 28]. Washington, DC is divided into eight Wards, and people who live in eastern and eastern-central Wards typically have fewer resources, less education, higher unemployment, lower income, and a higher rate of poor health indicators, such as obesity, diabetes, heart disease, and a shorter lifespan, than those in western and western-central Wards [27]. The population is also largely African American in the east and becomes predominantly Caucasian in the west. We predicted that demographic differences in Wards throughout Washington, DC would influence people’s knowledge and perceptions about the interconnectedness of natural resources, climate change, economics, and socio-cultural well-being, so we conducted surveys over three years to test that prediction. Understanding what people know and perceive, and which demographic characteristics may influence knowledge and perceptions, is key to designing effective educational programs, engaging in collective conversations, and building effective partnerships that find solutions for environmental problems and benefit the community.

## METHODS

The survey included five demographic questions (i.e., age, education, ethnicity, gender, and place of primary residence), and 26 open-ended, close-ended, and Likert scale questions/statements (hereafter “questions”) to assess knowledge and perceptions of the participants (Table 1, 2, 3). Some questions were duplicated or adapted from the connectedness-to-nature-scale [29]. We also included a question about whether the District government was spending the appropriate amount on health, workforce development, education, protecting natural resources, developing natural resources, law enforcement, and drug rehabilitation to see what people thought about spending for natural resources compared to other priority areas (Table 3). We loaded the survey into the iSurvey app (Harvest Your Data, Wellington, New Zealand) and trained undergraduate students in our classes at the University of the District of Columbia to conduct face-to-face interviews during each fall semester from 2016-2018. We canvassed 11 neighborhoods in Washington, DC’s eight Wards and solicited participants at transit stations, businesses, libraries, homes, and along sidewalks. Participants were adults (≥ 18 years old) and were selected because of their presence in the area only and without regard to any demographic category. In total, 455 completed surveys were collected. A survey was considered complete once a participant was read the final question, but it was not mandatory for the participants to answer every question.

**Table 1.** Open-ended questions that assessed knowledge and perceptions of people of different educational attainment in Washington, DC, USA about the interconnectedness of natural resources, climate change, economics, and socio-cultural well-being.

| Survey question | Education (sample size) | Responses (%) |  |  |  |  |  |  |  | Statistics |
| --- | --- | --- | --- | --- | --- | --- | --- | --- | --- | --- |
| Can you describe what natural resources are? | | Natural resources | Creation | Recycling | No | | | | | $\chi^2 = 15.4$<br>df = 6<br>$P = 0.018$ |
|  | Advanced degree (59) | 84.7 | 1.7 | 11.9 | 1.7 |  |  |  |  |  |
|  | College/trade (219) | 85.4 | 0.9 | 12.8 | 0.9 |  |  |  |  |  |
|  | High school (177) | 66.7 | 1.1 | 30.5 | 1.7 |  |  |  |  |  |
| What do you consider to be the most important natural resource? | | Air | Water | Soil | Trees | Land | Energy / fossil fuels | Multiple resources | Other | $\chi^2 = 18.8$<br>df = 14<br>$P = 0.17$ |
|  | Advanced degree (59) | 11.9 | 47.5 | 0 | 3.4 | 0 | 11.9 | 10.2 | 15.2 |  |
|  | College/trade (219) | 22.4 | 44.1 | 1.4 | 1.4 | 1.8 | 10.5 | 6.4 | 15.1 |  |
|  | High school (177) | 18.6 | 37.9 | 2.3 | 8.5 | 0 | 7.9 | 6.2 | 18.6 |  |
| Which natural resource has been threatened the most in your neighborhood? | | Air | Water | Soil | Trees | Land | Energy / fossil fuels | Multiple resources | Other | $\chi^2 = 15.6$<br>df = 14<br>$P = 0.34$ |
|  | Advanced degree (59) | 23.7 | 25.4 | 0 | 6.8 | 1.7 | 3.4 | 3.4 | 35.6 |  |
|  | College/trade (219) | 21.5 | 27.9 | 0 | 8.7 | 1.8 | 10.0 | 3.2 | 26.9 |  |
|  | High school (177) | 14.1 | 23.2 | 2.8 | 11.3 | 0.6 | 6.8 | 3.4 | 37.9 |  |
| Can you describe what climate change is to you? | | Weather patterns | Human cause / reaction | Climate change | No | | | | | $\chi^2 = 12.8$<br>df = 6<br>$P = 0.046$ |
|  | Advanced degree (59) | 66.1 | 15.3 | 11.9 | 6.8 |  |  |  |  |  |
|  | College/trade (219) | 61.6 | 12.3 | 18.3 | 7.8 |  |  |  |  |  |
|  | High school (177) | 64.4 | 9.0 | 8.5 | 18.1 |  |  |  |  |  |

**Table 2.**
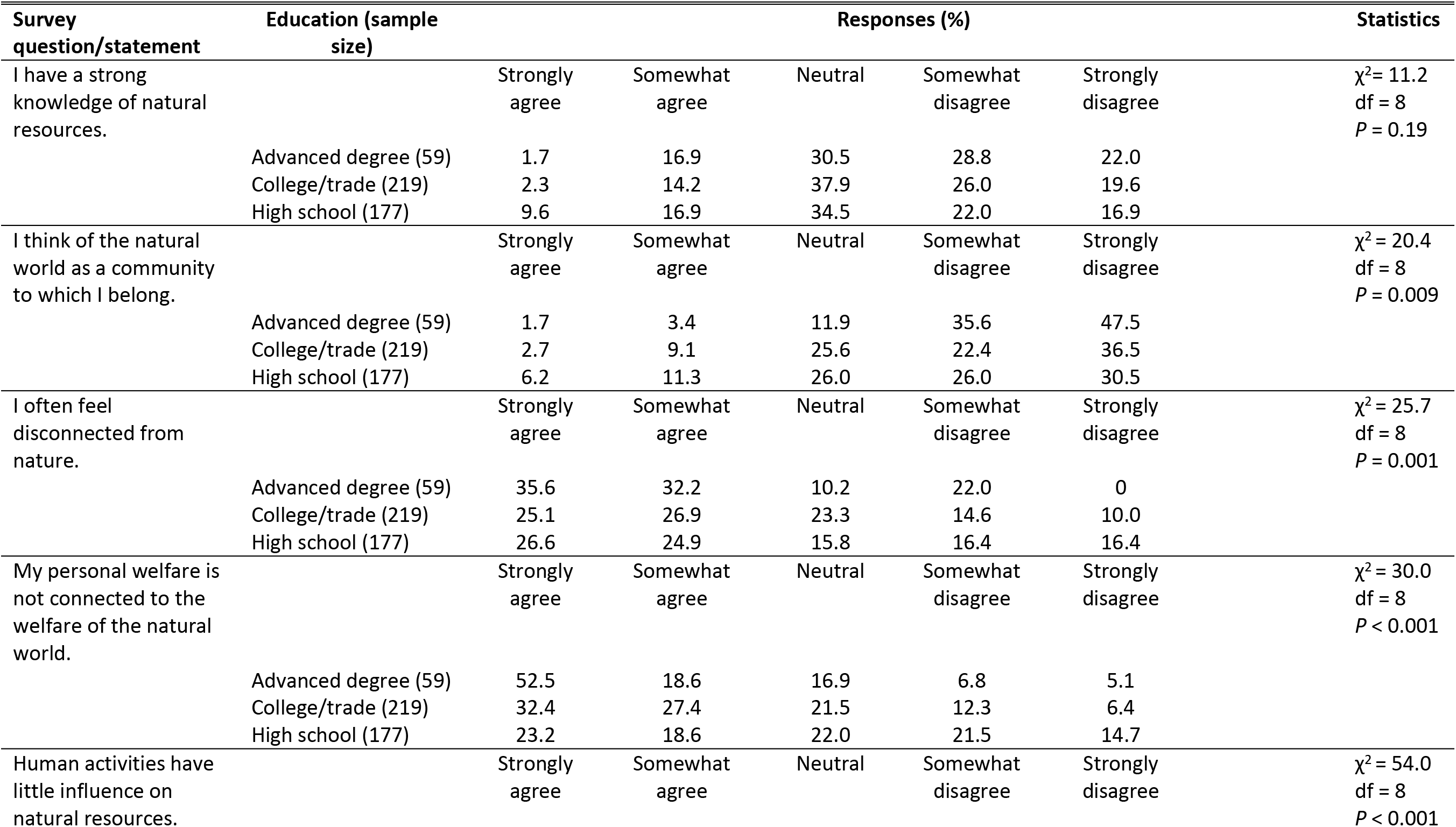
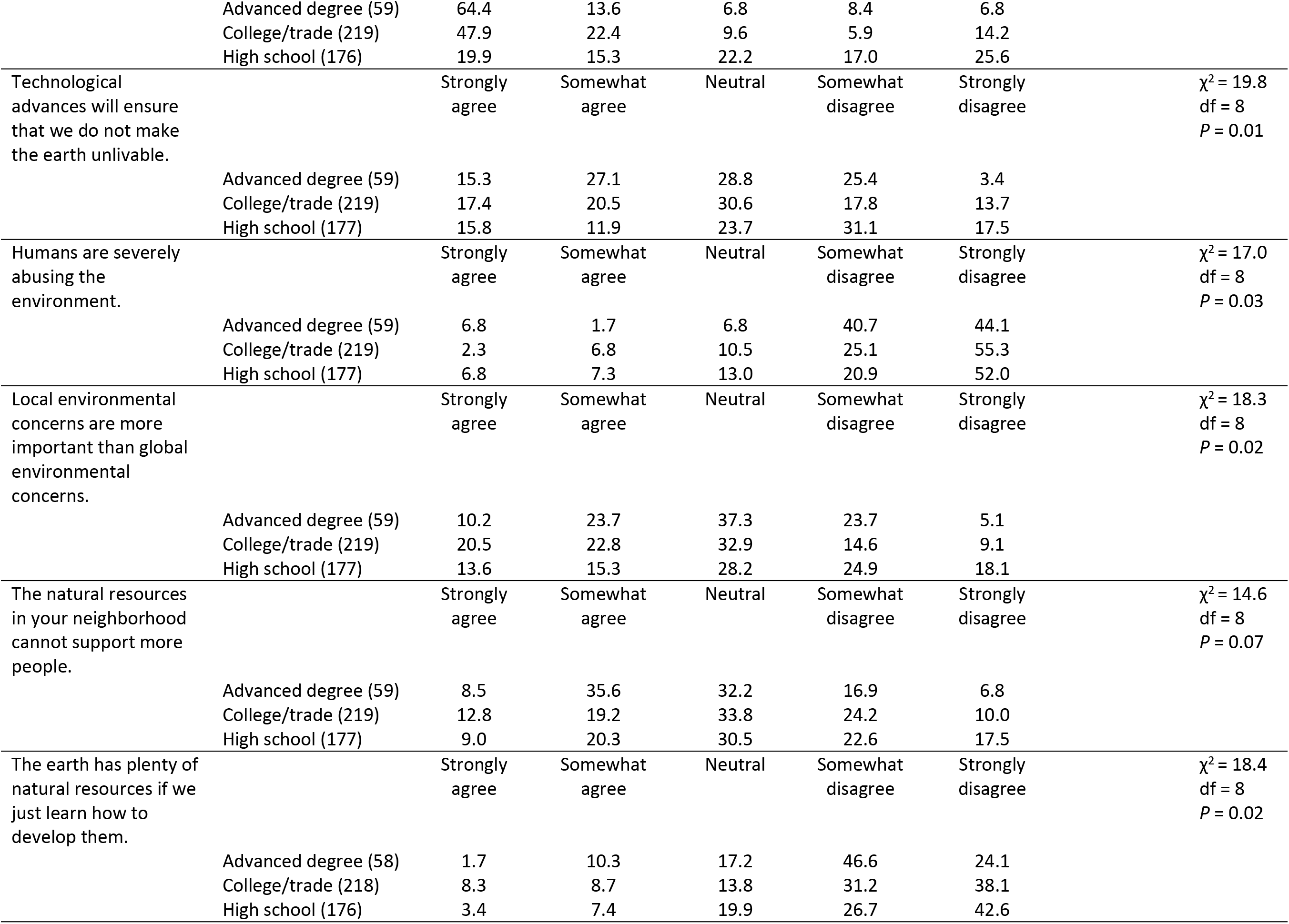
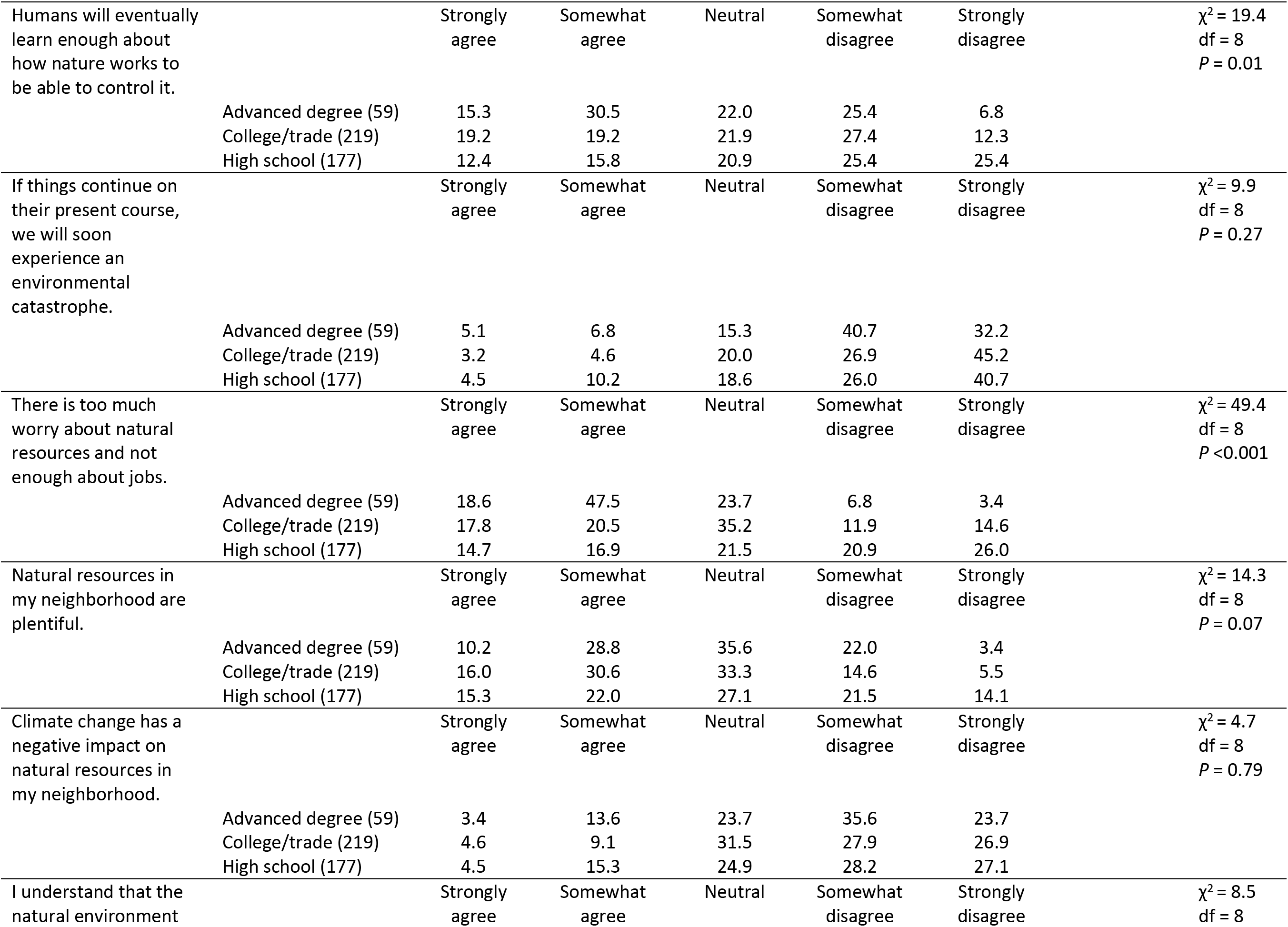
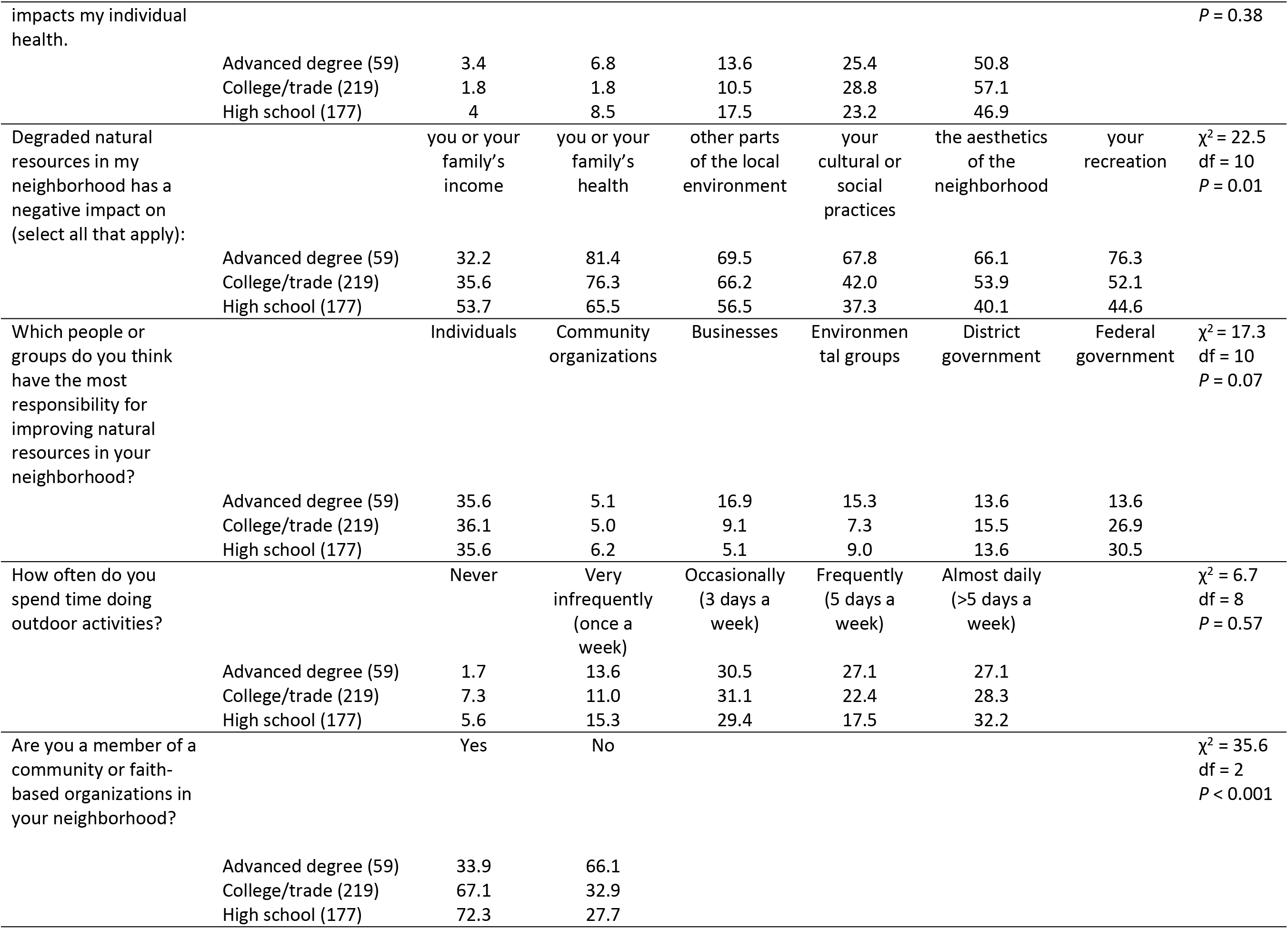
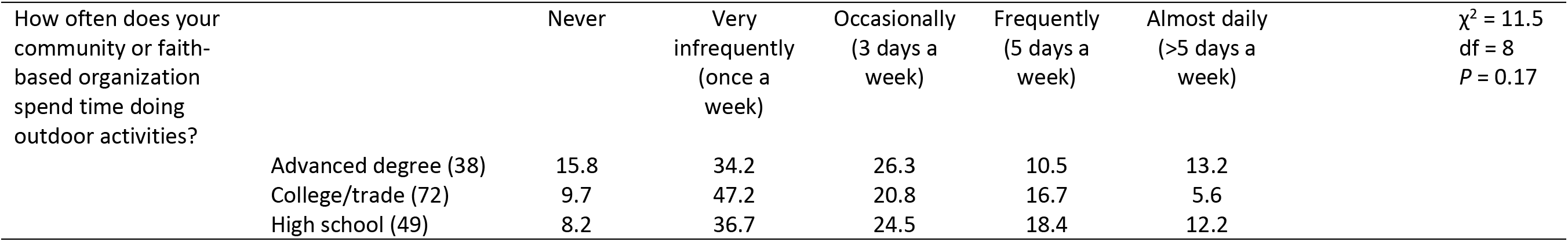
Close-ended and Likert scale questions/statements that assessed knowledge and perceptions of people of different educational attainment in Washington, DC, USA about the interconnectedness of natural resources, climate change, economics, and socio-cultural well-being.

| Survey question/statement | Education (sample size) | Responses (%) |  |  |  |  | Statistics |
| --- | --- | --- | --- | --- | --- | --- | --- |
| I have a strong knowledge of natural resources. | | Strongly agree | Somewhat agree | Neutral | Somewhat disagree | Strongly disagree | $\chi^2 = 11.2$<br>df = 8<br>$P = 0.19$ |
|  | Advanced degree (59) | 1.7 | 16.9 | 30.5 | 28.8 | 22.0 |  |
|  | College/trade (219) | 2.3 | 14.2 | 37.9 | 26.0 | 19.6 |  |
|  | High school (177) | 9.6 | 16.9 | 34.5 | 22.0 | 16.9 |  |
| I think of the natural world as a community to which I belong. | | Strongly agree | Somewhat agree | Neutral | Somewhat disagree | Strongly disagree | $\chi^2 = 20.4$<br>df = 8<br>$P = 0.009$ |
|  | Advanced degree (59) | 1.7 | 3.4 | 11.9 | 35.6 | 47.5 |  |
|  | College/trade (219) | 2.7 | 9.1 | 25.6 | 22.4 | 36.5 |  |
|  | High school (177) | 6.2 | 11.3 | 26.0 | 26.0 | 30.5 |  |
| I often feel disconnected from nature. | | Strongly agree | Somewhat agree | Neutral | Somewhat disagree | Strongly disagree | $\chi^2 = 25.7$<br>df = 8<br>$P = 0.001$ |
|  | Advanced degree (59) | 35.6 | 32.2 | 10.2 | 22.0 | 0 |  |
|  | College/trade (219) | 25.1 | 26.9 | 23.3 | 14.6 | 10.0 |  |
|  | High school (177) | 26.6 | 24.9 | 15.8 | 16.4 | 16.4 |  |
| My personal welfare is not connected to the welfare of the natural world. | | Strongly agree | Somewhat agree | Neutral | Somewhat disagree | Strongly disagree | $\chi^2 = 30.0$<br>df = 8<br>$P < 0.001$ |
|  | Advanced degree (59) | 52.5 | 18.6 | 16.9 | 6.8 | 5.1 |  |
|  | College/trade (219) | 32.4 | 27.4 | 21.5 | 12.3 | 6.4 |  |
|  | High school (177) | 23.2 | 18.6 | 22.0 | 21.5 | 14.7 |  |
| Human activities have little influence on natural resources. | | Strongly agree | Somewhat agree | Neutral | Somewhat disagree | Strongly disagree | $\chi^2 = 54.0$<br>df = 8<br>$P < 0.001$ |
|  | Advanced degree (59) | 64.4 | 13.6 | 6.8 | 8.4 | 6.8 |  |
|  | College/trade (219) | 47.9 | 22.4 | 9.6 | 5.9 | 14.2 |  |
|  | High school (176) | 19.9 | 15.3 | 22.2 | 17.0 | 25.6 |  |
| Technological advances will ensure that we do not make the earth unlivable. | | Strongly agree | Somewhat agree | Neutral | Somewhat disagree | Strongly disagree | $\chi^2 = 19.8$<br>df = 8<br>P = 0.01 |
|  | Advanced degree (59) | 15.3 | 27.1 | 28.8 | 25.4 | 3.4 |  |
|  | College/trade (219) | 17.4 | 20.5 | 30.6 | 17.8 | 13.7 |  |
|  | High school (177) | 15.8 | 11.9 | 23.7 | 31.1 | 17.5 |  |
| Humans are severely abusing the environment. | | Strongly agree | Somewhat agree | Neutral | Somewhat disagree | Strongly disagree | $\chi^2 = 17.0$<br>df = 8<br>P = 0.03 |
|  | Advanced degree (59) | 6.8 | 1.7 | 6.8 | 40.7 | 44.1 |  |
|  | College/trade (219) | 2.3 | 6.8 | 10.5 | 25.1 | 55.3 |  |
|  | High school (177) | 6.8 | 7.3 | 13.0 | 20.9 | 52.0 |  |
| Local environmental concerns are more important than global environmental concerns. | | Strongly agree | Somewhat agree | Neutral | Somewhat disagree | Strongly disagree | $\chi^2 = 18.3$<br>df = 8<br>P = 0.02 |
|  | Advanced degree (59) | 10.2 | 23.7 | 37.3 | 23.7 | 5.1 |  |
|  | College/trade (219) | 20.5 | 22.8 | 32.9 | 14.6 | 9.1 |  |
|  | High school (177) | 13.6 | 15.3 | 28.2 | 24.9 | 18.1 |  |
| The natural resources in your neighborhood cannot support more people. | | Strongly agree | Somewhat agree | Neutral | Somewhat disagree | Strongly disagree | $\chi^2 = 14.6$<br>df = 8<br>P = 0.07 |
|  | Advanced degree (59) | 8.5 | 35.6 | 32.2 | 16.9 | 6.8 |  |
|  | College/trade (219) | 12.8 | 19.2 | 33.8 | 24.2 | 10.0 |  |
|  | High school (177) | 9.0 | 20.3 | 30.5 | 22.6 | 17.5 |  |
| The earth has plenty of natural resources if we just learn how to develop them. | | Strongly agree | Somewhat agree | Neutral | Somewhat disagree | Strongly disagree | $\chi^2 = 18.4$<br>df = 8<br>P = 0.02 |
|  | Advanced degree (58) | 1.7 | 10.3 | 17.2 | 46.6 | 24.1 |  |
|  | College/trade (218) | 8.3 | 8.7 | 13.8 | 31.2 | 38.1 |  |
|  | High school (176) | 3.4 | 7.4 | 19.9 | 26.7 | 42.6 |  |
| Humans will eventually learn enough about how nature works to be able to control it. | | Strongly agree | Somewhat agree | Neutral | Somewhat disagree | Strongly disagree | $\chi^2 = 19.4$<br>df = 8<br>$P = 0.01$ |
|  | Advanced degree (59) | 15.3 | 30.5 | 22.0 | 25.4 | 6.8 |  |
|  | College/trade (219) | 19.2 | 19.2 | 21.9 | 27.4 | 12.3 |  |
|  | High school (177) | 12.4 | 15.8 | 20.9 | 25.4 | 25.4 |  |
| If things continue on their present course, we will soon experience an environmental catastrophe. | | Strongly agree | Somewhat agree | Neutral | Somewhat disagree | Strongly disagree | $\chi^2 = 9.9$<br>df = 8<br>$P = 0.27$ |
|  | Advanced degree (59) | 5.1 | 6.8 | 15.3 | 40.7 | 32.2 |  |
|  | College/trade (219) | 3.2 | 4.6 | 20.0 | 26.9 | 45.2 |  |
|  | High school (177) | 4.5 | 10.2 | 18.6 | 26.0 | 40.7 |  |
| There is too much worry about natural resources and not enough about jobs. | | Strongly agree | Somewhat agree | Neutral | Somewhat disagree | Strongly disagree | $\chi^2 = 49.4$<br>df = 8<br>$P < 0.001$ |
|  | Advanced degree (59) | 18.6 | 47.5 | 23.7 | 6.8 | 3.4 |  |
|  | College/trade (219) | 17.8 | 20.5 | 35.2 | 11.9 | 14.6 |  |
|  | High school (177) | 14.7 | 16.9 | 21.5 | 20.9 | 26.0 |  |
| Natural resources in my neighborhood are plentiful. | | Strongly agree | Somewhat agree | Neutral | Somewhat disagree | Strongly disagree | $\chi^2 = 14.3$<br>df = 8<br>$P = 0.07$ |
|  | Advanced degree (59) | 10.2 | 28.8 | 35.6 | 22.0 | 3.4 |  |
|  | College/trade (219) | 16.0 | 30.6 | 33.3 | 14.6 | 5.5 |  |
|  | High school (177) | 15.3 | 22.0 | 27.1 | 21.5 | 14.1 |  |
| Climate change has a negative impact on natural resources in my neighborhood. | | Strongly agree | Somewhat agree | Neutral | Somewhat disagree | Strongly disagree | $\chi^2 = 4.7$<br>df = 8<br>$P = 0.79$ |
|  | Advanced degree (59) | 3.4 | 13.6 | 23.7 | 35.6 | 23.7 |  |
|  | College/trade (219) | 4.6 | 9.1 | 31.5 | 27.9 | 26.9 |  |
|  | High school (177) | 4.5 | 15.3 | 24.9 | 28.2 | 27.1 |  |
| I understand that the natural environment | | Strongly agree | Somewhat agree | Neutral | Somewhat disagree | Strongly disagree | $\chi^2 = 8.5$<br>df = 8 |

|  |  |  |  |  |  |  |  |  |
| --- | --- | --- | --- | --- | --- | --- | --- | --- |
| impacts my individual health. | | | | | | | | $P = 0.38$ |
|  | Advanced degree (59) | 3.4 | 6.8 | 13.6 | 25.4 | 50.8 |  |  |
|  | College/trade (219) | 1.8 | 1.8 | 10.5 | 28.8 | 57.1 |  |  |
|  | High school (177) | 4 | 8.5 | 17.5 | 23.2 | 46.9 |  |  |
| Degraded natural resources in my neighborhood has a negative impact on (select all that apply): | | you or your family's income | you or your family's health | other parts of the local environment | your cultural or social practices | the aesthetics of the neighborhood | your recreation | $\chi^2 = 22.5$<br>df = 10<br>$P = 0.01$ |
|  | Advanced degree (59) | 32.2 | 81.4 | 69.5 | 67.8 | 66.1 | 76.3 |  |
|  | College/trade (219) | 35.6 | 76.3 | 66.2 | 42.0 | 53.9 | 52.1 |  |
|  | High school (177) | 53.7 | 65.5 | 56.5 | 37.3 | 40.1 | 44.6 |  |
| Which people or groups do you think have the most responsibility for improving natural resources in your neighborhood? | | Individuals | Community organizations | Businesses | Environmental groups | District government | Federal government | $\chi^2 = 17.3$<br>df = 10<br>$P = 0.07$ |
|  | Advanced degree (59) | 35.6 | 5.1 | 16.9 | 15.3 | 13.6 | 13.6 |  |
|  | College/trade (219) | 36.1 | 5.0 | 9.1 | 7.3 | 15.5 | 26.9 |  |
|  | High school (177) | 35.6 | 6.2 | 5.1 | 9.0 | 13.6 | 30.5 |  |
| How often do you spend time doing outdoor activities? | | Never | Very infrequently (once a week) | Occasionally (3 days a week) | Frequently (5 days a week) | Almost daily (>5 days a week) | | $\chi^2 = 6.7$<br>df = 8<br>$P = 0.57$ |
|  | Advanced degree (59) | 1.7 | 13.6 | 30.5 | 27.1 | 27.1 |  |  |
|  | College/trade (219) | 7.3 | 11.0 | 31.1 | 22.4 | 28.3 |  |  |
|  | High school (177) | 5.6 | 15.3 | 29.4 | 17.5 | 32.2 |  |  |
| Are you a member of a community or faith-based organizations in your neighborhood? | | Yes | No | | | | | $\chi^2 = 35.6$<br>df = 2<br>$P < 0.001$ |
|  | Advanced degree (59) | 33.9 | 66.1 |  |  |  |  |  |
|  | College/trade (219) | 67.1 | 32.9 |  |  |  |  |  |
|  | High school (177) | 72.3 | 27.7 |  |  |  |  |  |

| How often does your community or faith-based organization spend time doing outdoor activities? | | Never | Very infrequently (once a week) | Occasionally (3 days a week) | Frequently (5 days a week) | Almost daily (>5 days a week) | $\chi^2 = 11.5$<br>df = 8<br>$P = 0.17$ |
| --- | --- | --- | --- | --- | --- | --- | --- |
|  | Advanced degree (38) | 15.8 | 34.2 | 26.3 | 10.5 | 13.2 |  |
|  | College/trade (72) | 9.7 | 47.2 | 20.8 | 16.7 | 5.6 |  |
|  | High school (49) | 8.2 | 36.7 | 24.5 | 18.4 | 12.2 |  |

**Table 3.** Responses of people of different educational attainment in Washington, DC, USA when asked “For each of the following categories, answer whether you think the District government is spending too little, about the right amount, too much, undecided.”

| Category | Education (sample size) | Responses (%) |  |  |  | Statistics |
| --- | --- | --- | --- | --- | --- | --- |
|  |  | Too little | About the right amount | Too much | Undecided |  |
| Health | Advanced degree (59) | 54.2 | 30.5 | 5.1 | 10.2 | $\chi^2 = 2.6$ |
|  | College/trade (219) | 58.9 | 26.5 | 2.7 | 11.9 | df = 6 |
| | High school (177) | 62.1 | 26.5 | 2.7 | 11.9 | $P = 0.85$ |
| Workforce development | Advanced degree (59) | 49.2 | 33.9 | 5.1 | 11.9 | $\chi^2 = 1.7$ |
|  | College/trade (219) | 52.5 | 34.2 | 3.7 | 9.6 | df = 6 |
| | High school (177) | 54.2 | 28.8 | 6.2 | 10.7 | $P = 0.95$ |
| Education | Advanced degree (59) | 69.5 | 22.0 | 1.7 | 6.8 | $\chi^2 = 3.3$ |
|  | College/trade (219) | 64.4 | 26.0 | 1.8 | 7.8 | df = 6 |
| | High school (177) | 61.6 | 27.7 | 4.5 | 6.2 | $P = 0.78$ |
| Protecting natural resources | Advanced degree (59) | 57.6 | 25.4 | 0 | 16.9 | $\chi^2 = 6.5$ |
|  | College/trade (219) | 58.4 | 23.3 | 4.1 | 14.2 | df = 6 |
| | High school (177) | 54.2 | 26.0 | 6.2 | 13.6 | $P = 0.37$ |
| Developing natural resources | Advanced degree (59) | 57.6 | 27.1 | 1.7 | 13.6 | $\chi^2 = 9.3$ |
|  | College/trade (219) | 55.7 | 22.4 | 5.0 | 16.9 | df = 6 |
| | High school (177) | 46.9 | 26.6 | 10.7 | 15.8 | $P = 0.16$ |
| Law enforcement | Advanced degree (59) | 28.8 | 30.5 | 23.7 | 16.9 | $\chi^2 = 6.7$ |
|  | College/trade (219) | 30.6 | 26.5 | 32.9 | 10.0 | df = 6 |
| | High school (177) | 32.8 | 25.4 | 33.9 | 7.9 | $P = 0.35$ |
| Drug rehabilitation | Advanced degree (59) | 52.5 | 25.4 | 5.1 | 16.9 | $\chi^2 = 3.1$ |
|  | College/trade (219) | 55.7 | 20.1 | 6.8 | 17.4 | df = 6 |
| | High school (177) | 55.4 | 22.6 | 9.6 | 12.4 | $P = 0.79$ |

Some demographic variables were potentially correlated (e.g., people in some Wards are also likely from a certain ethnic group), so they were not all independent variables. Therefore, we ran preliminary analyses using separate χ^2^ contingency tests to determine whether the responses to two questions differed according to age, education, ethnicity, gender, and place of primary residence. We selected education as the focal demographic category because preliminary analyses found strong differences in response to the statement that “human activities have little influence on natural resources” (*P* < 0.05), whereas no strong differences in responses were found across other demographic variables (*P* > 0.05).

Participants were categorized into one of three groups based on their level of educational attainment: 1) completion of high school or less (hereafter “high school”) (n = 177); 2) some trade school or university education beyond high school up to and including completion of a trade school, two-, or four-year degree (hereafter “post-high school”) (219); and 3) completion of a Master’s, professional, or doctoral degree (hereafter “advanced education”) (n = 59).

There were four open-ended questions that probed participants’ perceptions of natural resources (Table 1). We coded answers to each open-ended question in order to reduce all responses to a limited number of categories. These response categories were analyzed using χ^2^ contingency tests to determine whether the responses differed among educational groups. Answers to the question “Can you describe what natural resources are?” were coded to fit into four categories: natural resources, creation, recycling, and none (Table 1). When the participant gave an example of a natural resource, such as air, water, trees, land, this indicated that they understood what natural resources are and their response was coded as “natural resources.” Participants’ answers that included terms such as God or biblical phrases were coded as “creation.” Answers that indicated a reuse or recycling of materials for financial gain were coded as “recycling.” Finally, responses that indicated that the participants were unable to answer the question were coded as “no.” Answers to the questions “What do you consider to be the most important natural resource?” and “Which natural resources has been threatened the most in your neighborhood?” were coded to fit into eight categories: air, water, soil, trees, land, energy/fossil fuels, multiple resources, and other (Table 1). Participants’ answers that included oil, fossil fuel, coal, and gas were coded as “energy/fossil fuels.” Participants’ answers that included more than one natural resource were coded as “multiple resources.” The “other” category includes natural resources that were infrequently mentioned, such as food, and resources that were not natural, such as education and transportation. Answers to the question “Can you describe what climate change is to you?” were coded to fit into four categories: weather patterns, human cause/reaction, climate change, and no.

Participants’ answers that included weather change, temperature change, hotter or colder weather, or similar statements were coded as “weather patterns.” Participants’ answers that voiced a human cause or invoked a human emotion, such as scary or fear, were coded as “human cause/reaction.” Participants’ answers that included climate change causes or effects (not including weather), such as global warming, carbon emissions, greenhouse gases, and sea level rise, were coded as “climate change.” Finally, responses that indicated that the participants were unable to answer the question were coded as “no.”

We also used separate χ^2^ contingency tests to determine whether the responses to the other 22 questions differed among educational groups. Sample size was sometimes fewer than the total number of participants previously reported when a participant did not answer a question.

## RESULTS

Participants across the three educational groups answered 14 questions dissimilarly (see Tables 1 and 2 for data supporting the results in this paragraph). Over 84% of participants in the post-high school and advanced education groups were able to describe natural resources, whereas fewer than 67% of participants in the high school group were able to do so. Participants in the high school group were more likely to discuss recycling of materials for financial gain when asked to describe natural resources. Over 60% of all participants discussed weather patterns when asked to describe climate change, but those with advanced degrees or post-high school education also discussed other causes and effects of climate change, whereas a greater percentage of those in the high school group were unable to describe climate change. Over 80% of participants with advanced education somewhat or strongly agreed that they think of the natural world as a community to which they belong, but only slightly over half of the other participants agreed with this question. Similarly, those with advanced education were less likely to feel disconnected from nature than other participants. Whereas over 70% of those with advanced education somewhat or strongly disagreed that their personal welfare is not connected to the welfare of the natural world, fewer than 60% and 42% of participants in the post-high school and high school groups, respectively, felt similarly. A relatively low percentage of participants with an advanced (15.2%) or post-high school education (20.1%) somewhat or strongly agreed that human activities have little influence on natural resources, whereas 42.6% of participants in the high school group somewhat or strongly agreed with this question. Those with the lowest educational attainment were also most likely to somewhat or strongly agree that technological advances will ensure that we do not make the earth unlivable (48.6% of the high school group versus 28.8-31.5% of other participants). Those with a post-high school or high school education were more likely to strongly agree that humans are severely abusing the environment than those with an advanced education, but 84.8% of those with an advanced education selected that they somewhat or strongly agreed with this question compared to 72.9-80.4% of other participants. Participants in the high school group were more likely to somewhat or strongly agree that local environmental concerns are more important than global concerns (43% versus 23.7-28.8% of participants with post-high school and advanced education). Participants with a post-high school or high school education were more likely than those with an advanced education to strongly agree that earth has plenty of natural resources if we just learn how to develop them. Those in the high school group were also more likely than other participants to strongly agree that humans will eventually learn enough about how nature works to be able to control it and to somewhat or strongly agree that there is there is too much worry about natural resources and not enough about jobs. Those with an advanced education were over twice as likely to be a member of a community organization or faith-based group than all other participants.

Participants were asked whether degraded natural resources in their neighborhood had a negative impact on income, health, environment, cultural/social practices, neighborhood aesthetics, and recreation (see Table 2 for data supporting the results in this paragraph). More than two-thirds of participants with an advanced education selected that local degraded natural resources negatively impacted all of these except income. Income was selected by fewer than a third of participants with an advanced education. Those in the post-high school group felt less strongly than those in the advanced education group that degraded natural resources had an impact on these categories; however, more than half still thought that local degraded natural resources negatively impacted health, environment, neighborhood aesthetics, and recreation. Participants in the high school group felt less strongly than all other participants that local degraded natural resources impact these categories, except for income. More than half of these participants said that degraded natural resources negatively impacted income. Health and the environment were the other two categories where more than half of those in the high school group said that local degraded natural resources had a negative impact.

Participants across all educational groups answered 12 questions similarly (see Tables 1 and 2 for data supporting the results in this paragraph). All participants most frequently mentioned water as the most important natural resource and the one most threatened in their neighborhood, followed by air. Participants were more likely to agree that they had a strong knowledge of natural resources than disagree, but the most common answer was “neutral.” Participants frequently selected “neutral” to the statement that natural resources in their neighborhood cannot support more people, with participants with advanced education somewhat disagreeing with this statement and all others somewhat agreeing. Participants also selected “neutral” most frequently to the statement that natural resources in their neighborhood are plentiful. Over 66% of participants somewhat or strongly agreed that if things continue on their present course, we will soon experience an environmental catastrophe, over 55% somewhat or strongly agreed that climate change negatively impacts natural resources in their neighborhood, and over 70% somewhat or strongly agreed that they understand that the natural environment impacts their health. Over one-third of participants thought individuals had the most responsibility to improve natural resources, followed by government entities (federal or district government). Businesses, environmental groups, and community organizations were less frequently selected. Approximately half the participants reported spending time outdoors frequently (5 days per week) or almost daily, although those that belong to a faith-based or community organization said their organization was outdoors infrequently (once per week).

Participants were also in agreement about spending by Washington, DC’s government and most frequently thought the government was spending “too little” on each of the seven priority areas (Table 3). All participants were especially likely to say that District government spends too little on education (>60% of participants). Those in the high school group were also especially likely to say that too little was spent on health (62.1% of participants). Over 54% of all participants thought District government spent too little on protecting natural resources and over 46% thought too little was spent on developing natural resources. Law enforcement was the priority area for which participants were least likely to say that spending was too little.

## DISCUSSION

People in Washington, DC had some similar knowledge and perceptions about the interconnectedness of natural resources, climate change, economics, and socio-cultural well-being. Whereas survey participants did not report having a strong knowledge about natural resources, most were able to define natural resources, listed water and air as the natural resources that they were most concerned about, indicated that the natural environment affects health, and reported that climate change negatively impacts natural resources. What created similar knowledge and perceptions among participants is unknown, but could be due to shared experiences, such as lived experiences and exposure to these issues through education and the media, and/or shared values. In fact, nationally in the USA there has been an increased awareness and concern about at least one major environmental issue: climate change [30]. A national survey about climate change found that people are increasingly discussing it with family and friends, regularly exposed to it in the media, and reporting that they feel the effects of climate change and are harmed by them [30]. The participants in the national survey also expressed worry about extreme weather events, especially those pertaining to water, such as flooding, drought, and shortages [30]. Some of the similarities among participants in our survey in Washington, DC may be part of the shifting attitudes and knowledge happening on a national scale. Additionally, the “biospheric (concern for environment)” and “altruistic (concern for others and intrinsic value)” value orientations influence responses to environmental issues and climate change [31], so the people in Washington, DC may have similar values.

Despite some similarities among survey participants, educational attainment, as we predicted, influenced people’s knowledge and perceptions about the interconnectedness of natural resources, climate change, economics, and socio-cultural well-being. Whereas most participants could describe natural resources, a large percentage of those in the high school group could not. People with an advanced education showed a greater understanding of climate change and its impacts, which is consistent with a global survey that found that educational attainment was the strongest predictor of awareness about climate change [32]. Those with an advanced degree were also most likely to report that their personal welfare depends on the natural community and reported the highest connection with the natural community. Connection to nature is often correlated with time spent outdoors [33], but we found that time spent outdoors was similar across educational groups. We speculate that the type of activities engaged in outdoors are more likely to result in a greater feeling of connectedness-to-nature than the amount of time. We base this speculation on the fact that those with an advanced degree were most likely to report that degraded natural resources impacted their recreation, which may indicate that outdoor activities are more commonly recreational with this educational group compared to other groups.

People in the high school group were most likely to believe that humans do not have much influence on natural resources and placed more trust in technology and human achievements to control nature and ensure that earth will not become unlivable; beliefs that are not uncommon [34], but likely incorrect without concurrent changes to population growth and resource exploitation [35]. Compared to those with education beyond high school, those with a high school education were also most likely to express an interest in local environmental concerns over global, jobs over natural resources, and effects of degraded local natural resources on income, health, and the environment instead of on cultural/social practices, neighborhood aesthetics, and recreation. Education is correlated with employment and income, with unemployment declining and income increasing with educational attainment [36], which may explain why those with no education beyond high school are more concerned about the local environment and its impact on jobs, income, and health. Vulnerable and marginalized people, such as those who are undereducated, poor, in a minority racial or ethnic group, and/or an immigrant are also disproportionately afflicted by climate change, a degraded environment, and environmental hazards [37–40], so they may be more acutely aware of the local environment and its effects on prosperity and well-being.

People afflicted by poor ecosystem health and degradation of natural resources need educational opportunities, tools of empowerment to change their circumstances, and employment that affords them to the choice to relocate or adapt, such as jobs in the clean energy sector. Results from the survey suggest topics that could be emphasized through formal educational classes, cooperative extension programs, traditional media, social media, and other platforms in order to increase knowledge about environmental issues and their interrelationship with economics and socio-cultural well-being. For example, understanding of natural resources is lower than climate change among all survey participants and fewer than half the participants reported having a strong knowledge of natural resources, so natural resources could be emphasized. Understanding of environmental issues, the influence of people on natural resources, and the connection between the natural world and their personal welfare is lower in people who have not had schooling beyond high school. Since as many as one-quarter of students graduating from high school in the USA are completing a science curriculum that is below standard [41], the deficiencies in knowledge may stem, in part, from lack of exposure in primary and secondary school. The deficiencies could be addressed by providing opportunities for education. However, we found that some environmental knowledge was relatively high even among the less educated group, which is consistent with previous studies that show high awareness of environmental risks and support for environmental protections regardless of education and across racial groups that may, on average, be less educated than Caucasians [42–45].

Beyond education that creates awareness of environmental issues, people from diverse groups must be given tools of empowerment that enable them to change their circumstances and demonstrate pro-environmental actions, such as advocating for environmental policies. One way in which to empower people is to recruit them into active roles in group organizations. Globally, environmental knowledge and action are often correlated with participation in group organizations [12] and participants in our study with an advanced degree generally had high environmental knowledge and reported higher involvement in community or faith-based organizations. Ensuring that people with lower educational attainment have equal opportunities to participate in group organizations may help close the gap in environmental knowledge, provide a tool of empowerment to change their circumstances and take pro-environmental actions. Mainstream environmental organizations have a low percentage of non-white minorities on their staff [46] and the term “environmentalist” is associated with well-educated and white people by minorities and Caucasians alike [45], so structural and psychological barriers currently prevent diverse participation and representation.

Our results suggest that those wishing to lead collective conversations and build effective partnerships that find solutions for environmental problems need to take the demographics of their stakeholders into account. Stakeholders with advanced degrees may be likely to think and act more globally and show more of an interest in curtailing environmental problems that have a negative impact on their recreation, neighborhood aesthetics, and cultural/social practices. However, engaging stakeholders with a high school education means shifting focus to local concerns and issues that have a more immediate impact on their jobs and income. People with lower incomes will more likely want to discuss mitigation and adaptation measures in their neighborhoods, such as improved emergency alerts, access to government subsidies for air conditioners and energy-efficient appliances, stronger buildings that withstand extreme weather, and more local agriculture and community gardens [47]. The negative impacts of a degraded environment on health and the environment, especially air and water, are common concerns that would likely be of interest to most of the population in Washington, DC, regardless of their educational attainment. So, for example, increasing awareness among the population in Washington, DC that degradation of the environment promotes poor air quality, which exacerbates chronic illnesses, may prompt people to want to more thoroughly understand natural resources and climate change. The survey participants indicated that individuals, followed by government entities, have the greatest responsibility to improve local natural resources, so people should be empowered to engage in the process of improving a degraded environment and taught how to advocate for changes within the government.

## ACKNOWLEDGMENTS

We thank the hundreds of survey participants, the dozens of students who helped interview survey participants, and Caitlin Arlotta, Jennifer Bennett-Mintz, Jenifer Gonzalez, and Elizabeth Jewett for facilitating some interview sessions. This project was partially funded by USDA Joint Venture Agreement 15-JV-11242306-076 between the USDA Northeast Climate Hub and the University of the District of Columbia.

## REFERENCES

1. United Nations. World urbanization prospects. Department of Economic and Social Affairs. Population Division. 2014 [cited 2019 September 11] Available from: http://www.un.org/en/development/desa/news/population/world-urbanization-prospects-2014.html

2. United States Census Bureau. Growth in urban population outpaces rest of nation. U.S. Department of Commerce, U. S. Census Bureau, Washington, DC, USA. 2012 [cited 2019 September 11] Available from: https://www.census.gov/newsroom/releases/archives/2010_census/cb12-50.html

3. Gleeson T, Wada Y, Bierkens MFP, van Beek LPH. Water balance of global aquifers revealed by groundwater footprint. Nature. 2012; 488: 197–200.

4. Ye Y, Gutierrez NL. Ending fishery overexploitation by expanding from local successes to globalized solutions. Nat Ecol Evol. 2017; 1:0179.

5. Dirzo R, Young HS, Galetti M, Ceballos G, Isaac NJ, Collen B. Defaunation in the Anthropocene. Science. 2014; 345: 401–406.

6. Hallmann CA, Sorg M, Jongejans E, Siepel H, Hofland N, Schwan H, Stenmans W, Müller A, Sumser H, Hörren T, Goulson D, de Kroon H. More than 75 percent decline over 27 years in total flying insect biomass in protected areas. PLOS ONE. 2017; 12:e0185809.

7. Stocker TF, Qin D, Plattner G-K, Tignor M, Allen SK, Boschung J, Nauels A, Xia Y, Bex V, Midgley PM, editors. Climate change 2013: the physical science basis. Contribution of working group I to the fifth assessment report of the Intergovernmental Panel on Climate Change. Cambridge: Cambridge University Press; 2014.

8. McDonald RI. Conservation for cities. How to plan and build natural infrastructure. Washington, DC: Island Press; 2015.

9. Romm J. Climate change: what everyone needs to know. New York: Oxford; 2016.

10. Kollmuss A, Agyeman J. Mind the gap: why do people act environmentally and what are the barriers to pro-environmental behavior. Environ Educ Res. 2002; 8: 239–260.

11. Lorenzoni I, Nicholson-Cole S, Whitmarsh L. Barriers perceived to engaging with climate change among the UK public and their policy implications. Global Environ. Change. 2007; 17: 445–459.

12. Shih-Wu L, Wei-Ta F, Shin-Cheng Y, Shiang-Yao L, Tsai H, Chou J-Y, Ng E. A nationwide survey evaluating the environmental literacy of undergraduate students in Taiwan. Sustainability. 2018; 10:1730.

13. Satterthwaite D, McGranahan G, Tacoli C. Urbanization and its implications for food and farming. Philosophical Transactions of the Royal Society B Biological Sciences. 2010; 365: 2809–2820.

14. Yigitcanlar T, Dizdaroglu D. Ecological approaches in planning for sustainable cities: a review of the literature. Global J Environ Sci Manag. 2015; 1:71–94.

15. Sun S, Tian L, Cao W, Lai P-C, Wong PPY, Lee RS-y, Mason TG, Krämer A, Wong C-M. Urban climate modified short-term association of air pollution with pneumonia mortality in Hong Kong. Sci Tot Environ. 2019; 646: 618–624.

16. Larson LR, Jennings V, Cloutier SA. Public parks and wellbeing in urban areas of the United States. PLOS ONE. 2016; 11:e0153211.

17. Erickson LE, Jennings M. Energy, transportation, air quality, climate change, health nexus: sustainable energy is good for our health. AIMS Public Health. 2017; 4: 47–61.

18. Erickson LE, Griswold W, Maghirang RG, Urbaszewski BP. Air quality, health and community action. J Environ Prot. 2017; 8: 1057–1074.

19. Carlson K, McCormick S. American adaptation: social factors affecting new developments to address climate change. Glob. Environ. Chang. 2015; 35: 360–367.

20. District of Columbia Department of Energy and the Environment. Plans and commitments. 2019 [cited 11 September 19]. Available from: https://doee.dc.gov/service/plans-and-commitments

21. District of Columbia Department of Energy and the Environment. Climate ready DC. 2019 [cited 11 September 19]. Available from: https://doee.dc.gov/climateready

22. Kalkstein LS, Sailor D, Shickman K, Sheridan S, Vanos J. Assessing the health impacts of urban heat island reduction strategies in the District of Columbia. 2013 [cited 2019 September 11]. Available from: https://doee.dc.gov/sites/default/files/dc/sites/ddoe/publication/attachments/20131021_Urban%20Heat%20Island%20Study_FINAL.pdf

23. McCright AM. The effects of gender on climate change knowledge and concern in the American public. Popul Environ. 2010; 32: 66–87.

24. Ogunbode CA, Arnold K. A study of environmental awareness and attitudes in Ibadan, Nigeria. Hum Ecol Risk Assess: An Int J. 2012; 18: 669–684.

25. Stevenson KT, Peterson MN, Bondell HD, Mertig AG, Moore SE. Environmental, institutional, and demographic predictors of environmental literacy among middle school children. PLOS ONE. 2013; 8:e59519.

26. Rajapaksa D, Islam M, Managi S. Pro-environmental behavior: The role of public perception in infrastructure and the social factors for sustainable development. Sustainability 2018; 10: 937.

27. District of Columbia Department of Health. Health equity report: District of Columbia. 2018. [cited 11 September 19]. Available from: https://app.box.com/s/yspij8v81cxqyebl7gj3uifjumb7ufsw

28. Turner MA. Poor people and poor neighborhoods in the Washington metropolitan area. Washington, DC: Urban Institute; 1997.

29. Mayer FS, McPherson-Frantz I. The connectedness to nature scale: a measure of individuals’ feeling in community with nature. J Environ Psychol. 2004; 24:503–515.

30. Leiserowitz A, Maibach E, Rosenthal S, Kotcher J, Ballew M, Goldberg M, Gustafson A. Climate change in the American mind: December 2018. Yale University and George Mason University. New Haven, CT: Yale Program on Climate Change Communication. 2018 [cited 2019 September 11] Available from: https://climatecommunication.yale.edu/publications/climate-change-in-the-american-mind-december-2018/2/

31. Marshall NA, Thiault L, Beeden A, Beeden R, Benham C, Curnock MI, Diedrich A, Gurney GG, Jones L, Marshall PA, Nakamura N, Pert P. (2019 Our environmental value orientations influence how we respond to climate change. Front Psychol. 2019; 10:1–8.

32. Lee TM, Markowitz EM, Howe PD, Ko C-Y, Leiserowitz AA. Predictors of public climate change awareness and risk perception around the world. Nat Clim Change. 2015; 5: 1014–1020.

33. Larson LR, Szczytko R, Bowers EP, Stephens LE, Stevenson KT, Floyd MF. Outdoor time, screen time, and connection to nature: troubling trends among rural youth? Environ Behav. 2018; 1–26.

34. Ausubel JH. Can technology spare the earth? Amer. Sci. 1996; 84: 166–178.

35. Huesemann MH, Huesemann JA. Will progress in science and technology avert or accelerate global collapse? A critical analysis and policy recommendations. Environ Dev Sustain. 2008; 10: 787–825.

36. Wolla SA, Sullivan J. Education, income, and wealth. Saint Louis: Page One Economics; 2017. Available from: https://files.stlouisfed.org/files/htdocs/publications/page1-econ/2017-01-03/education-income-and-wealth_SE.pdf

37. Congressional Black Caucus Foundation. African Americans and climate change: an unequal burden. Redefining Progress, Washington, DC.

38. Ebi KL, Balbus J, Kinney PL, Lipp E, Mills D, O’Neill MS, Wilson M. Effects of global change on human health. In: Gamble, JL, lead author. Analyses of the Effects of Global Change on Human Health and Welfare and Human Systems. Washington, DC: Synthesis and Assessment Product 4.6. U.S. Environmental Protection Agency; 2008. pp. 39–87.

39. Karl TR, Melillo JM, Peterson TC (eds). Global climate change impacts in the United States. Cambridge: Cambridge University Press, Cambridge; 2009.

40. Mohai P, Pellow D, Roberts JT. Environmental justice. Annu Rev Environ Resour. 2009; 34:405–430.

41. Nord C, Roey S, Perkins R, Lyons M, Lemanski N, Brown J, Schuknecht J. The Nation’s Report Card: America’s High School Graduates (NCES 2011-462). Washington, D.C.: U.S. Department of Education, National Center for Education Statistics; 2011.

42. Dietz T, Dan A, Shwom R. Support for climate change policy: Social psychological and social structural influences. Rural Sociol. 2007; 72: 185–214.

43. Leiserowitz A, Akerlof K. Race, ethnicity and public responses to climate change. New Haven, CT Yale Program on Climate Change Communication. 2010) [cited 2019 September 25] Available from: Available at climatecommunication.yale.edu/publications/ race-ethnicity-and-public-responses-to-climate-change/. Accessed October 16, 2018.

44. Macias T. Environmental risk perception among race and ethnic groups in the United States. Ethnicities. 2016; 16: 111–129.

45. Pearson AR, Schuldt JP, Romero-Canyas R, Ballew MT, Larson-Konar D. Diverse segments of the US public underestimate the environmental concerns of minority and low-income Americans. Proc Natl Acad Sci USA. 2018; 115: 12429–12434.

46. Taylor, DE. The state of diversity in environmental organizations: Mainstream NGOs, foundations, government agencies. 2014 [cited 2019 September 25] Available from: http://www.diversegreen.org/the-challenge/.

47. Kreslake JM. Perceived importance of climate change adaptation and mitigation according to social and medical factors among residents of impacted communities in the United States. Health Equity 2019; 3: 124–133.

